# HaloTrace: A spatiotemporally precise fluorescent readout of blood-brain barrier permeability in mice

**DOI:** 10.1101/2025.10.30.685531

**Authors:** Hannah L. Zucker, Ekaterina Konshina, Perry Mitchell, Chenghua Gu

## Abstract

The blood-brain barrier (BBB) is an indispensable, selectively permeable interface that controls the entry and exit of nutrients, ions and waste products into the brain. Despite its biological importance, most measurements of BBB permeability rely on dyes that suffer from nonspecific signals, lack of spatial fidelity, and incompatibility with longitudinal or repeated measurements. Here we present HaloTrace: a method which leverages the HaloTag ligand-receptor tool to generate a precise spatiotemporal readout of BBB integrity that avoids major pitfalls of existing methods. We present evidence that the fluorescent HaloTag ligand has minimal interactions with blood contents but can enter the brain specifically at sites of BBB dysfunction, where it covalently binds to nearby HaloTag receptors. The ligand accumulates in the brain during its short lifetime in circulation and is stably anchored in place for at least 24 hours. Unlike existing tracers, free ligand is not retained in the blood vessels at detectable levels, so the entirety of ligand fluorescence represents true BBB leakage. Furthermore, we demonstrate that HaloTrace can quantify BBB permeability at multiple discrete timepoints prior to the experiment endpoint. This offers researchers the ability to study the progression or resolution of BBB permeability in a way current methods cannot. HaloTrace is thus uniquely poised to characterize the spatiotemporal dynamics of BBB leakage in mouse models.

## INTRODUCTION

The blood-brain barrier (BBB), formed by endothelial cells that comprise the walls of central nervous system (CNS) blood vessels, performs an essential role in maintaining CNS homeostasis by restricting the movement of substances into and out of the CNS.^1^ The structural bases for the BBB’s restrictiveness are the presence of tight junctions that seal the spaces between adjacent brain endothelial cells and the active suppression of transcytosis, which is the trafficking of substances across the endothelium through the endocytic network.^1^ Key molecular players in these structures have been identified, including the essential TJ component Claudin5^2–4^ and Mfsd2a, a membrane protein that suppresses caveolae formation and thus downregulates transcytosis at the BBB.^5,6^ Researchers have also identified signaling pathways that regulate BBB function, such as canonical Wnt signaling in ECs through β-catenin.^7–9^ However, much is still unknown about the molecular components and regulators of BBB function. Dynamics of essential processes, including tight junction assembly/maintenance,^4^ speed and routes of transcytosis,^6^ precise onset of BBB loss-of-function in disease models,^10,11^ and timescales of subsequent repair^12^ are areas of active investigation. If we are to uncover the molecular mechanisms that govern BBB permeability, and the timescales under which they operate, we need techniques capable of quantifying BBB dynamics. Yet, existing readouts of BBB permeability have relatively poor spatiotemporal resolution, can be difficult to quantify, and lack capacity to capture dynamic properties of BBB leakage across time.

Conventional measurements of BBB permeability, often referred to as leakage assays, involve injection of an exogenous molecule (i.e., tracer) into the bloodstream. In mice without BBB disruption, the tracer will be contained within the blood vessel lumen. However, in mice with increased BBB permeability, the tracer will leak across the BBB into the brain where it can be visualized as leakage hotspots. Thus, tracer presence in brain tissue indicates an aberrant increase in BBB permeability. A wide variety of tracers have been utilized, such as fluorophore or biotin conjugates for fluorescence microscopy, horseradish peroxidase for electron microscopy, or gadolinium contrast media for MRI.^2,6,9,13^

Popular tracers have enabled great insight into BBB research yet have significant caveats: (1) they interact with biological molecules in blood, (2) they have low spatiotemporal specificity, (3) they cannot be used to make multiple measurements across time in a single mouse, and (4) they are readily detectable in blood during circulation and after histological sample preparation, which makes observing modest BBB leakage difficult or impossible. Some popular tracers even have biological interactions that can compromise vascular health. For example, long-chain dextrans conjugated to fluorophores can elicit anaphylactic reactions in some animal models, and high doses of Horseradish Peroxidase have shown deleterious side effects.^13^

Many tracers, such as Evans Blue and sulfo-NHS-biotin, bind to proteins in the blood and brain.^14,15^ Even sulfo-NHS-biotin, a common modern tracer, readily reacts with free amines on proteins at physiological pH via its N-hydroxysuccinimide moiety. This reactivity means that depending on the concentration injected, common tracers circulate in both free and bound forms in proportion to the total tracer concentration in circulation. Thus, it is impossible to interpret whether the BBB is permissive to small, unbound tracer molecules, larger tracer-protein complexes, or both. Apart from the implications for overall BBB permeability, tracer size is also diagnostic of the type of BBB leakage. Tight junction dysfunction (paracellular leakage) is thought to be detected only with small tracers, whereas dysregulation of transcytosis (transcellular leakage) can be detected with small and large tracers.^2^ Thus, the information that biologically reactive tracers can tell us about a BBB permeability phenotype is inherently limited.

Furthermore, tracers that leak into the brain can be spatially redistributed or washed away by brain fluid dynamics or sample preparation.^16^ Because tracer in the brain does not always remain fixed at the site of BBB leakage, existing tracer assays can only report BBB leakage accurately at one timepoint shortly after the tracer is injected into circulation. So, while they can report the presence and intensity of BBB leakage, existing methods cannot be used to determine exactly from which area the tracer leaked across the BBB or track leakage phenotypes over time. Finally, most fluorescent tracers are used in high concentrations that result in appreciable brightness in blood vessels which must be accounted for during data processing.

A more informative BBB tracer would have minimal interactions with blood contents yet remain anchored in the brain at locations where BBB integrity is lost. It would also be capable of comparing leakage intensity and spatial distribution at multiple timepoints within a single animal. To this end, we designed a two-component method for visualizing BBB leakage called HaloTrace in which the tracer is a small fluorescent ligand for a brain-localized HaloTag receptor.^17^ Using a well-established mouse model of BBB dysfunction, we show HaloTrace produces a quantifiable readout of the spatial extent of BBB leakage during multiple short timepoints in a single mouse.

## RESULTS

### HaloTrace design and characterization

HaloTrace is based on the HaloTag ligand-receptor system.^17^ The HaloTag system consists of two parts: a receptor protein (HaloTag) and a fluorescent ligand that together form a covalent bond (**Figure 1A**). The ligand, a haloalkane, has high receptor affinity and no known interactions with mammalian systems.^17^ Similarly, the HaloTag receptor is not endogenous to mammalian systems so its expression must be induced ectopically in cell types of interest. Importantly, components of HaloTrace have no reported biological interactions like those often observed for existing tracers.^17^

**Figure 1.**
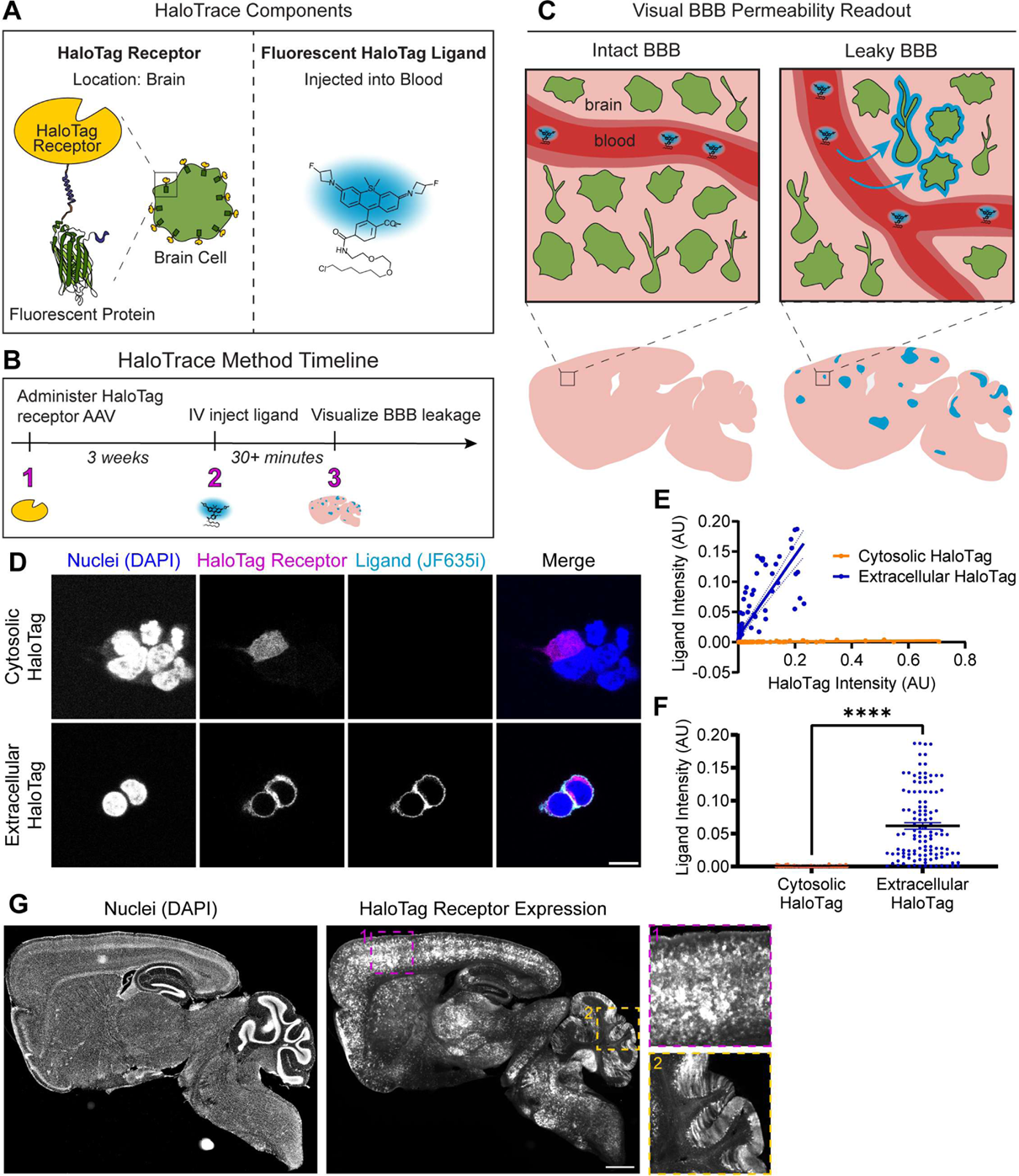
HaloTrace design and characterization. **(a)** The two components of HaloTrace are HaloTag receptor localized to the extracellular membrane of brain cells and its fluorescent ligand injected into the systemic blood circulation. **(b)** To perform the HaloTrace leakage assay, HaloTag receptor protein is expressed on the extracellular surface of brain cells using AAV-PHP.eB. Later, a cell-impermeable fluorescent ligand is injected into the bloodstream. Post mortem, brain sections are imaged to determine location and abundance of ligand in the brain. **(c)** The ligand should be excluded from the brain by the intact BBB but will leak into the brain if the BBB is compromised. Any ligand that leaks across the BBB rapidly covalently binds to the extracellular HaloTag receptors. The bound ligand, which is resistant to washout and anchored in place for the lifetime of the HaloTag receptor, can be visualized with confocal microscopy. **(d)** *In vitro* characterization in HEK cells of the extracellular HaloTag receptor localization and JF635i-ligand binding ability compared to a cytosolic HaloTag control. Merged image shows HaloTag JF635i-ligand (cyan), receptor (mScarlet, magenta), and nuclei (DAPI, blue). Scale bar: 10 µm. **(e)** Relationship between HaloTag receptor expression (mScarlet fluorescence) and JF635i ligand fluorescence intensity in extracellular HaloTag and cytosolic HaloTag conditions. Data points represent single cells transfected with extracellular HaloTag (blue, n=123) or cytosolic HaloTag (orange, n=112). Linear regression for extracellular HaloTag: y=0.6681x+0.009312, r^2^=0.68. Linear regression for cytosolic HaloTag: y=0.003822x+ 0.0001079, r^2^=0.45. **(f)** Quantification of JF635i ligand fluorescence intensity in extracellular HaloTag (blue) and cytosolic HaloTag (orange) cells. Data are mean ± SEM, extracellular n = 111, cytosolic n = 120 cells, ****p< 0.0001, Welch’s unpaired t test. **(g)** Representative slide scanner image of extracellular HaloTag receptor expression visualized with mScarlet (gray) achieved by systemic injection of AAV-PHP.eB-CAG- HaloTagTM-mScarlet at 2e11 vg dose. Nuclei are visualized with DAPI (blue). Scale bar: 1 mm. Insets highlight HaloTag receptor location in the cortex (dotted blue lines) and cerebellum (dotted yellow lines).

We envisioned a BBB permeability assay in which HaloTag receptors are expressed on the surface of CNS cells and a fluorescent cell-impermeable HaloTag ligand is injected into the bloodstream (**Figure 1B**). At sites of BBB leakage, ligand will enter the brain and forms a covalent bond with a HaloTag receptor, thereby fluorescently labeling locations in the brain where BBB integrity is lost (**Figure 1C)**. The fluorescent readout of BBB leakage is predicted to be stable for multiple days because any ligand covalently bound to a receptor will remain in place for the lifetime of the receptor.^18^ In contrast, previous work has demonstrated that similar fluorescent HaloTag ligands are rapidly eliminated from systemic circulation, with most ligand gone within ∼1h when injected at 100 nmol dose.^18,19^ Free ligand is not retained in the blood vessels at detectable levels, so the entirety of ligand fluorescence represents true BBB leakage. This rapid ligand clearance also produces the added benefit of restricting the measurement window for HaloTrace corresponding to the time period when ligand is present in circulation. Like other recently published HaloTag-based reporters of physiological processes, the measurement time is gated on the presence of free ligand, so the ligand distribution in brain is a fluorescent record of BBB leakage during its ∼1h lifetime in blood circulation.^18,20^ Therefore, the HaloTrace assay should produce a tight temporal readout of BBB permeability in the form of a long-lasting signal that can be visualized long after the ligand is gone from systemic circulation.

We first designed a DNA construct to express HaloTag receptor on the extracellular membrane (**Supplemental Figure 1A)**. We included targeting motifs to direct the HaloTag receptor to the extracellular face of the cell membrane.^21,22^ We also included the fluorescent protein mScarlet fused to the internal side of the transmembrane domain to visualize the HaloTag receptor location.^23^

For the HaloTag ligand, we chose fluorescent JF635i-ligand and JF549i-ligand because both are cell-impermeable, and thus are not predicted to cross the intact BBB.^19,24^ JF635i in particular is fluorogenic, meaning that its fluorescence increases dramatically when receptor-bound.^19^ Its fluorogenicity, combined with its short half-life in blood and easy washout of free ligand, ensure the ligand fluorescence observed in the HaloTrace assay corresponds to true leakage, not a nonspecific signal. At 825 Da and 747 Da for JF635i-ligand and JF549i-ligand, respectively, both should be small enough to detect paracellular leakage between junctions.^2^

We first performed an *in vitro* characterization of the extracellular HaloTag receptor and ligand. We transfected human embryonic kidney cells with the extracellular HaloTag receptor-mScarlet or a cytosolic HaloTag fused to mScarlet, then introduced JF635i-ligand into the cells’ media. The extracellular HaloTag receptor shows the expected localization to the extracellular membrane and successfully binds the JF635i-ligand (**Figure 1D**). In contrast, the cytosolic HaloTag receptor shows no detectable JF635i-ligand binding, indicating the JF635i-ligand is indeed cell-impermeable. The extracellular HaloTag receptor levels (indicated by mScarlet fluorescence intensity) correlate positively with JF635i ligand intensity, whereas in the control condition JF635i intensity does not increase even at the highest cytosolic HaloTag receptor expression (**Figure 1E**). Quantified another way, extracellular HaloTag expressing cells have statistically significantly higher total JF635i ligand intensity than cytosolic HaloTag expressing cells (**Figure 1F**). These results demonstrate that the extracellular HaloTag construct is correctly targeted to the extracellular face of the cell membrane, the JF635i-ligand is cell-impermeable, and the receptor-ligand interaction is highly specific with low background signal. The same is true when the ligand is left in the cell media for 24 hours **(Supplementary Figure 1B,C,D)**. We also observed specifically extracellular binding of the red cell-impermeable JF549i-ligand to an extracellular HaloTag fused to green fluorescent protein (GFP) rather than mScarlet (Supplementary Figure 1E,F).

After verifying the HaloTrace assay functioned as expected *in vitro*, we translated it into an *in vivo* setting. To accomplish this, we expressed the HaloTag receptor in the CNS by systemic administration of the AAV capsid PHP.eB. This AAV transduces neurons, astrocytes and other CNS cell types (**Figure 1G**).^25^ HaloTag-expressing cells include putative neurons, astrocytes, and other cell types in accordance with the reported tropism of AAV-PHP.eB.^25^ Interestingly, we observed a lack of HaloTag receptor expression in the choroid plexus, a CNS region lacking a BBB **(Supplemental Figure 1G)**. Overall, the AAV-mediated HaloTag receptor expression is well-positioned to bind biologically relevant extravasated ligand.

### HaloTrace reveals BBB leakage without an intravascular signal

We next tested HaloTrace’s ability to detect BBB leakage by using a well-characterized genetic model of BBB disruption, endothelial-specific inducible knockdown of β-catenin (Cdh5:CreER/+; Ctnnb1^f/f^).^9,26^ Within days of tamoxifen-induced *Ctnnb1* knockdown, these mice have significant attenuation of Wnt signaling in endothelial cells and profound loss of BBB integrity especially in the molecular layer of the cerebellum. Previous work has demonstrated Cdh5:CreER/+; Ctnnb1^f/f^ mice have increased BBB permeability to a variety of small and large tracers.^9^

Using HaloTrace, we observed areas of cerebellum in Cdh5:CreER/+; Ctnnb1^f/f^ mice with significant ligand enrichment (**Figure 2A**). The Ctnnb1^f/f^ control mice had comparatively little ligand fluorescence. Therefore, HaloTrace can correctly distinguish between a leaky and intact BBB. In the Cdh5:CreER/+; Ctnnb1^f/f^ mice, the pattern of ligand deposition in the cerebellum overlaps with the more broad receptor expression, indicating successful and enduring ligand-receptor binding to a fraction of HaloTag receptors (**Figure 2A,D**). To quantify leakage with HaloTrace, we used a CellProfiler algorithm to identify the total area of the cerebellum, the area occupied by HaloTag receptor (HaloTag+ area), and the area occupied by the ligand (ligand+ area) in each image. Then we computed the ligand+ area as a percent of the total HaloTag+ area, which was statistically significantly higher in the knockout mice compared to controls (**Figure 2B**). Although there was some variability in the HaloTag+ area in the cerebellum across mice (**Figure 2C**), the BBB permeability readout of the HaloTrace method, ligand+ area, was robust to these small differences. We also confirmed that the 100 nmol ligand dose we chose was not receptor saturating; increasing the ligand to 200 nmol resulted in higher ligand labeling area and integrated intensity (**Supplementary Figure 2C-F**).

**Figure 2.**
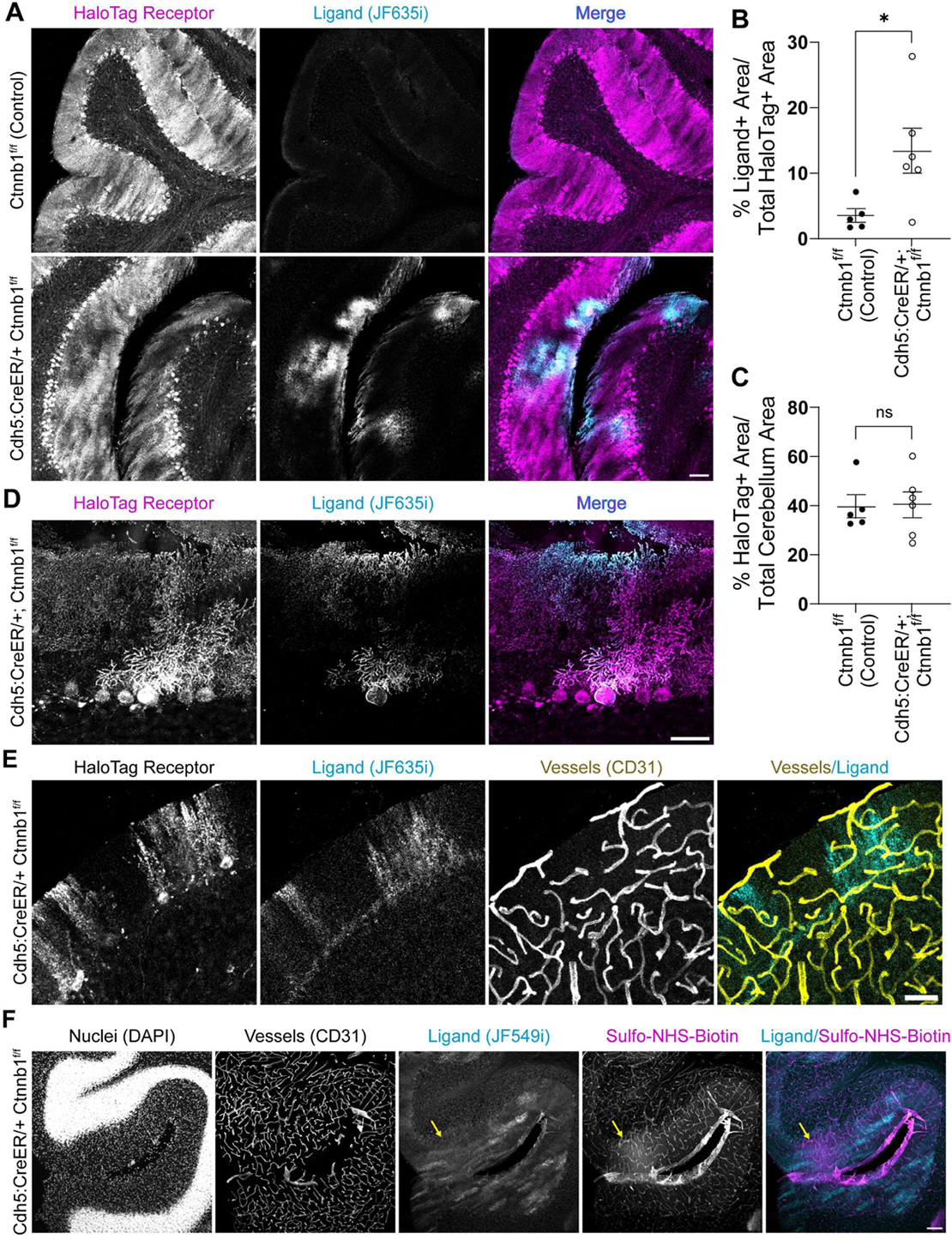
HaloTrace measures BBB leakage without an intravascular signal. **(a)** Representative 10x magnification confocal images of cerebellum show HaloTag receptor (magenta) and JF635i-ligand (cyan) after 0.75 h circulation of 100 nmol JF635i-ligand in Cdh5:CreER/+; Ctnnb1^f/f^ and Ctnnb1^f/f^ mice. Scale bar: 100 µm. **(b)** Quantification of ligand+ area as a percentage of HaloTag receptor expression area in cerebellum. Data represent mean ± SEM, Cre+ n=6, Cre- n=5, *p<0.05, nested t test. **(c)** Quantification of HaloTag positive area in cerebellum. Data represent mean ± SEM, Cre+ n=6, Cre- n=5, ns = not significant, nested t test. **(d)** Representative 40x magnification confocal images of leakage hotspots in Cdh5:CreER/+; Ctnnb1^f/f^ cerebellum. Scale bar: 50 µm. **(e)** Representative 10x magnification confocal image showing HaloTag receptor and JF635i-ligand (cyan) alongside vessels (CD31, yellow) in cerebellum of a Cdh5:CreER/+; Ctnnb1^f/f^ mouse. Scale bar: 50 µm. **(f)** Comparison of HaloTrace JF549i ligand (cyan) and sulfo-NHS-biotin tracer (magenta) localization in Cdh5:CreER/+; Ctnnb1^f/f^ cerebellum. Yellow arrow indicates an area of sulfo-NHS-biotin extravasation. Scale bar: 100 µm.

One drawback of existing methods is the retention of tracer inside vessels even after transcardial perfusion, so we next assessed whether HaloTrace suffers from the same phenomenon. Co-staining HaloTrace sections with a vessel marker shows no evidence of JF635i-ligand retention inside vessels (**Figure 2E**). Similarly, there is no appreciable intravascular ligand in brains that were processed without a perfusion fixation step to wash out the vascular contents (**Supplementary Figure 2A**). Although mild endothelial tropism has been reported for AAV-PHP.eB, we did not observe considerable receptor or ligand enrichment in endothelial cells.^25^

To compare the HaloTrace assay with a commonly used tracer assay, we performed HaloTrace and sulfo-NHS-biotin leakage assay simultaneously in Cdh5:CreER/+; Ctnnb1^f/f^ mice. We delivered HaloTag receptor AAV with a GFP tag, then induced gene knockdown with tamoxifen, and assayed permeability with a combination of JF549i ligand and sulfo-NHS-biotin. During immunofluorescence processing, we stained for a vessel marker (CD31). The data suggest, as previously established, that sulfo-NHS-biotin is retained in vessels even in areas without readily detectable tracer extravasation; in contrast, HaloTrace ligand is not observed inside vessels and does not need to be accounted for during quantification (**Figure 2F, Supplemental Figure 2B**). Overall, HaloTrace produced discrete, visualizable and quantifiable hotspots of leakage compared to the diffuse pattern produced by the sulfo-NHS-biotin leakage.

### HaloTrace records a stable snapshot of BBB leakage that persists for at least 24 hours

One limitation of BBB leakage assays is the requirement that histological samples must be collected soon after the tracer injection time. In contrast, we hypothesized that HaloTrace produces a stable, long-lasting signal that would allow greater flexibility in experimental design. To determine whether HaloTrace produced a stable fluorescent readout of BBB leakage independent of the tracer injection time, we next tested HaloTrace in a cohort of mice with the same conditions except we extended the delay between ligand injection and perfusion to 24 hours (**Figure 3A**). We hypothesized that the total amount of ligand in the brain would be equivalent in both time conditions because of the short lifetime of the ligand in circulation and the predicted longevity of receptor-bound ligand in the brain. The 24-hour delay group showed a comparable ligand enrichment pattern in the cerebellum of knockouts but not controls (**Figure 3B**). The ligand-positive area was again statistically significantly higher in knockouts than controls, as expected (**Figure 3C**). Total HaloTag area was again the same across genotypes, as expected (**Figure 3D**). The mean ligand positive area of the 24-hour group (13.40 ± 2.16 percent) was comparable to the 0.75-hour circulation condition (11.76 ± 1.87 percent). Importantly, the ligand is still restricted to locations with receptor expression after 24 hours (**Figure 3E**).

**Figure 3.**
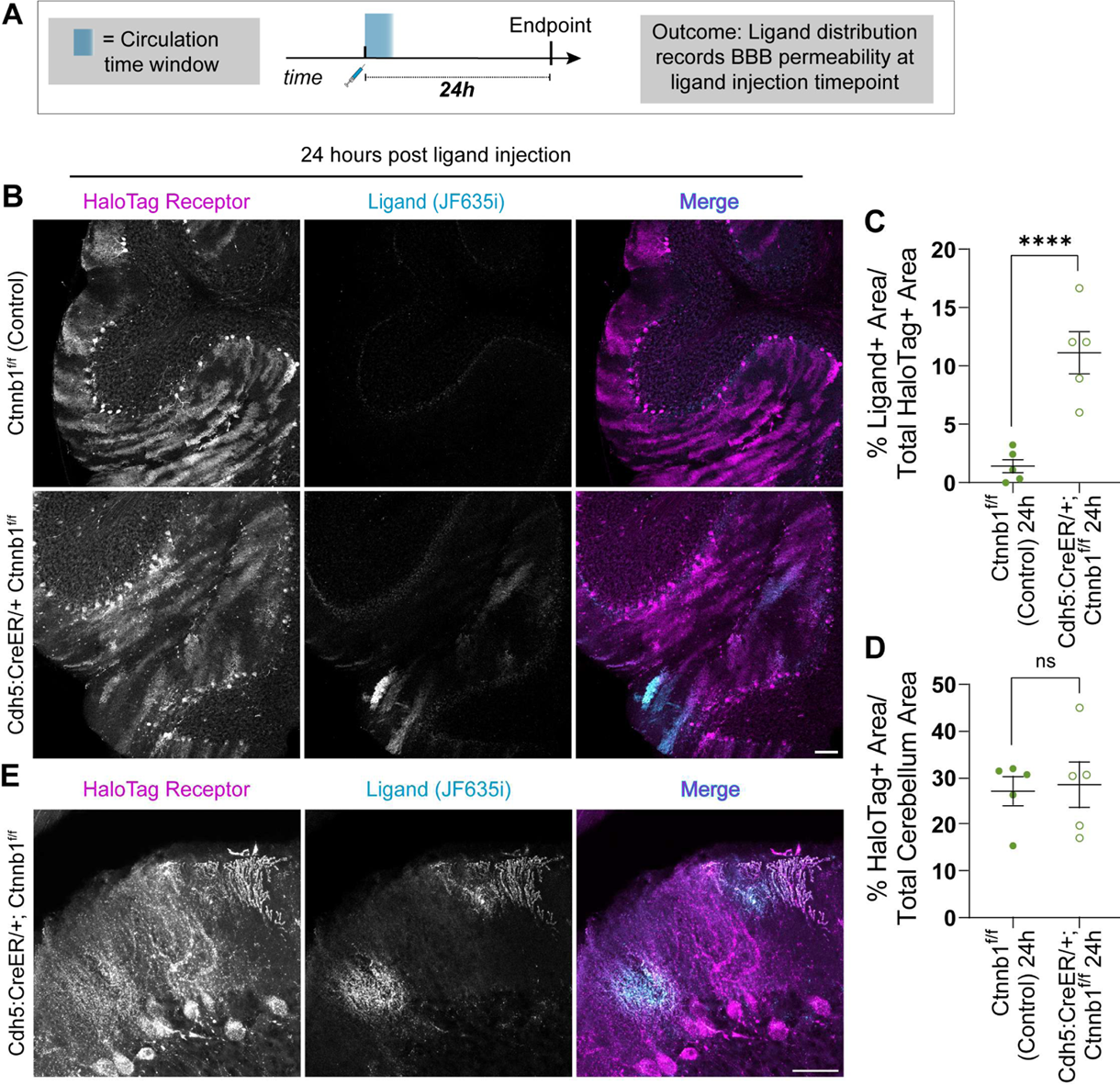
HaloTrace records a stable snapshot of BBB leakage that persists for at least 24 hours. **(a)** Diagram showing the experimental design. **(b)** Representative 10x magnification confocal images of cerebellum show HaloTag receptor (magenta) and JF635i-ligand (cyan) at 24 h following ligand circulation in Cdh5:CreER/+; Ctnnb1^f/f^ and Ctnnb1^f/f^ mice. Scale bar: 100 µm. **(c)** Quantification of ligand+ area as a percentage of HaloTag receptor expression area in cerebellum. Data represent mean ± SEM, n=5 per genotype, ****p<0.0001, nested t test. **(d)** Quantification of HaloTag positive area in cerebellum as a control. Data represent mean ± SEM, n=5 per genotype, ns = not significant, nested t test. **(e)** Representative 40x magnification confocal images of leakage hotspots at the 24 hour timepoint. Scale bar: 50 µm.

In summary, HaloTag receptor expression is similar across both genotypes, but the JF635i-ligand positive area is increased in knockdown mice. This indicates the ligand accumulation peaks in the time between ligand injection and experimental endpoint (0.75 hours), so HaloTrace reports leakage that occurred within 0.75 hours of ligand injection. Measurement of this readout can be delayed at least 24 hours. Although the experimental endpoint was delayed one day in the 24-hour group compared to the 0.75-hour group, both measurements faithfully correspond to a “snapshot” of leakage that occurred in the short window of time directly following ligand injection.

### HaloTrace records spatiotemporally specific leakage history at multiple timepoints

One of the greatest drawbacks of existing BBB assays is the difficulty of characterizing dynamic BBB phenotypes. For example, one may wonder whether the “hotspot” leakage pattern observed in the Cdh5:CreER/+; Ctnnb1^f/f^ cerebellum reflects static areas of BBB vulnerability or simply stochasticity in tracer distribution. We reasoned that HaloTrace’s unique combination of features – short tracer circulation time and longevity of the covalently anchored ligand – positions us to make sequential leakage measurements in the same mouse to address questions of this nature. As a proof of principle, we compared BBB leakage at two timepoints in the same Cdh5:CreER/+; Ctnnb1^f/f^ mouse using the spectrally compatible JF635i- and JF549i-ligands.^19^ In one group of mice, we injected the first ligand at 1.25 hours pre-perfusion fixation and the second at 0.75 hours, a 0.5h inter-injection interval (Δ0.5h). In the second group, we injected the ligands at 24 and 0.75 hours, respectively, a 24h interval (Δ24h) (**Figure 4A**). We chose these experimental timepoints because prior characterization with sulfo-NHS-biotin tracer indicates this model has similar BBB leakage intensity at days 8 and 9 post-tamoxifen.^9^ Then, we assessed the reliability of the dual timepoint readout by answering three important questions: (1) does JF549i-ligand perform similarly to the JF635i-ligand, indicating they are directly comparable? (2) does the order of the ligand administration or the delay between ligands change the leakage readout, meaning the ligands can be injected sequentially without interfering with each other? and (3) do we see less spatial overlap of the two ligands when we increase the interval between measurements, indicating a biologically relevant dynamic change in the spatial pattern of leakage across time?

**Figure 4.**
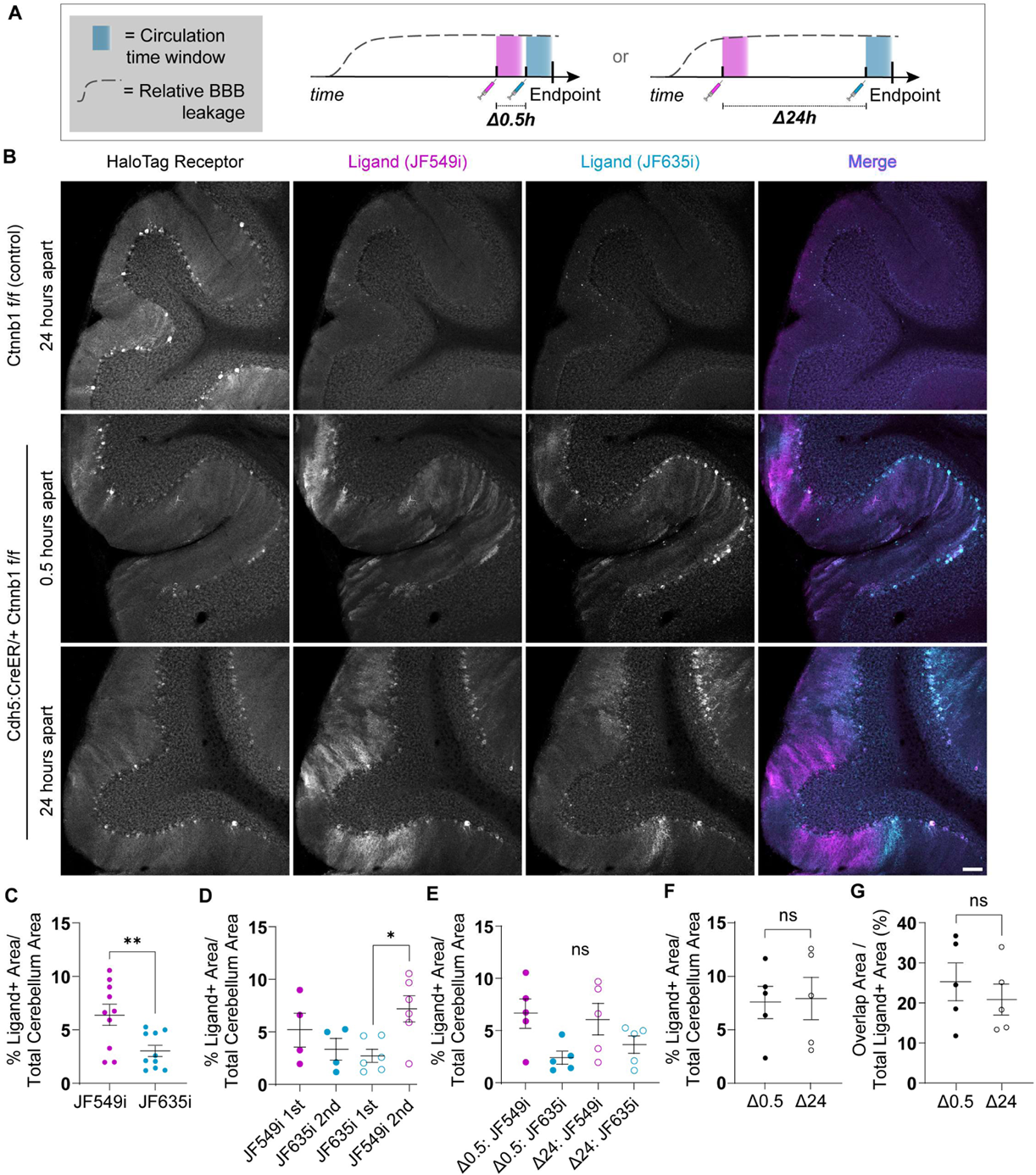
Sequential HaloTrace measurements capture BBB leakage patterns across time in a single mouse. **(a)** Experimental overview. Measurement time window shown in blue. Dotted line shows the relative BBB permeability over time. In the Δ0.5 group, one ligand is injected 1.25h prior to tissue fixation/collection and another is injected at 0.75h prior. In the Δ24 group, the first ligand is injected 24 hours prior to tissue collection and the second is injected at 0.75h prior. **(b)** Representative 10x magnification confocal images of the HaloTag receptor (gray), JF549i-ligand (magenta in merge), and JF549i-ligand (cyan in merge) in Ctnnb1^f/f^ and Cdh5:CreER/+; Ctnnb1^f/f^ cerebellum in the Δ0.5 hours and Δ24 hours cohorts. In the merge channel, areas of high ligand overlap appear as white. Scale bar: 100 µm. **(c)** Quantification of total JF549i-ligand and JF635i-ligand positive area across all conditions. Data represent mean ± SEM, n=10 mice, **p<0.01, nested t test. **(d)** Quantification of JF549i-ligand and JF635i-ligand positive area grouped by order of ligand administration. Data represent mean ± SEM, n= 4-6 mice per group, *p<0.05, unlabeled = non significant, nested ANOVA. **(e)** Quantification of JF549i-ligand and JF635i-ligand positive area grouped by the interval between ligand injections. Data represent mean ± SEM, n= 5 mice per group, *p<0.05, unlabeled = non significant, nested ANOVA. **(f)** Quantification of total ligand positive area in the Δ0.5 hours and Δ24 hours cohorts. Data represent mean ± SEM, n= 5 mice per group, ns = non significant, nested t test. **(g)** Quantification of the JF549i-ligand and JF635i-ligand overlap area as a percent of total ligand positive area in the Δ0.5 hours and Δ24 hours cohorts. Data represent mean ± SEM, n= 5 mice per group, ns = non significant, nested t test.

First we confirmed that administration of both ligands in sequence did not disrupt BBB integrity in a healthy Ctnnb1^f/f^ control (**Figure 4B**). In the Cdh5:CreER/+; Ctnnb1^f/f^ mice we observed that the two measurements revealed distinct but partially overlapping leakage hotspots in the cerebellum (**Figure 4B, Supplementary Figure 3A**). Across conditions, the JF549i-ligand hotspot areas were larger than corresponding JF635i-ligand hotspots, perhaps because of inherent differences in fluorophore properties (**Figure 4C**).

We next found that hotspot area for JF549i-ligand and JF635i-ligand did not vary depending on the order of administration (**Figure 4D**). However, the difference in intensity between JF549i- and JF635i-ligand was still apparent. Similarly, we found no evidence that ligand hotspot area varied based on the delay period between ligand injections, regardless of which ligand was injected first (**Figure 4E**).

Even though the JF549i-ligand occupied a greater area than the JF635i-ligand, this relationship was consistent across conditions and order of ligand administration, so it was still possible to compare the relationship between their leakage patterns across time. We found that the total area occupied by at least one ligand did not differ between the Δ0.5 hours and Δ24 hour groups (**Figure 4F**). We also found that the overlap of hotspot area was equivalent at Δ0.5 hours and Δ24 hours (**Figure 4G**). Intensity-based Mander’s and Pearson’s colocalization analyses corroborated the lack of difference between the Δ0.5 hours and Δ24 hour conditions (**Supplemental Figure 3B,C**). Together, these results confirm that HaloTrace can indeed perform sequential independent measurements of BBB permeability. Future work is required to determine whether the differences in leakage pattern produced by the two ligands were due to inherent differences in ligand properties, stochasticity in local BBB leakage, or a biologically relevant change in BBB permeability across time.

## DISCUSSION

Here we presented HaloTrace, a method to quantify BBB integrity in mouse models. We used the established Cdh5:CreER/+; Ctnnb1^f/f^ model of BBB dysfunction to showcase the useful and unique properties of HaloTrace, chiefly its temporospatial specificity lack of intravascular signal, stable and long-lived leakage signal, and capacity to report BBB leakage from multiple sequential timepoints in a single animal. Because quantification of HaloTrace is decoupled from its measurement time(s), it can be applied to a variety of experimental questions.

One advantageous feature of the HaloTrace method is its customizability. Its modular components enable one to customize the receptor location (e.g. on a specific cell type or brain region of interest) and the ligand properties (e.g., fluorescence, size, and chemical properties) to meet diverse experimental needs. Future work should characterize HaloTrace in other mouse models of BBB dysfunction, including those with exclusively tight junction defects. Previously, researchers have used a battery of tracers with divergent size, hydrophobicity, and solubility to assess tight junction permeability, but HaloTag ligands of different sizes but otherwise identical chemistry could cleanly reveal size-specific permeability features associated with tight junction defects.^4^

HaloTrace is not without downsides, but its customizability means potential workarounds for these drawbacks do exist. Its two-component design makes it more complicated to implement than single dye tracer assays. Importantly, the HaloTag receptor gene is delivered via AAV. The successful delivery of AAV-PHP.eB-HaloTag might be perturbed in mice with brain endothelial dysfunction. This can be avoided by administering the HaloTag AAV prior to the onset of BBB dysfunction, as we did in our experiments by injecting AAV prior to the induction of *Ctnnb1* knockdown and BBB leakage. This timing may not be possible or desirable for all applications. Alternatively, one could change the strategy for HaloTag receptor expression. We chose to deliver the HaloTag receptor with AAV because we could readily apply it in Cre-loxP mouse models, but another group has produced a mouse line with Tet-inducible cell-type specific expression of extracellular HaloTag that could be used in other experimental paradigms.^27^ An extracellular HaloTag mouse line could also prove useful for measuring leakage at early developmental timepoints.

Another constraint on the HaloTrace assay is the limited number of spectrally distinct fluorescent proteins and HaloTag ligands. The fluorescent protein marker in the HaloTag construct occupies a fluorescent channel that limits the ability to co-stain for multiple markers and image with commercially available confocal microscopes. To free more of the visual spectrum, HaloTrace could be demarcated with an epitope tag (i.e., V5 tag) rather than a fluorescent protein; verification of HaloTag expression could be achieved with immunofluorescence staining against the epitope tag. Freeing up an additional fluorescence channel would also make it possible to measure leakage at three timepoints with three different fluorescent ligands. Perhaps other fluorescent ligands in development are more similar than JF635i and JF549i and would thus produce directly comparable leakage readouts without further optimization. Commercially available fluorescent HaloTag ligands can be expensive, so it would also be useful to generate lower-cost alternatives.

HaloTrace’s key advantage over existing tracers is its capacity to compare the spatial distribution and extent of BBB leakage at multiple timepoints in the same experimental animal. Although we did not observe conclusive evidence of BBB permeability dynamics at the timepoints we tested in the *Ctnnb1* knockout mouse, the question remains: why do so many BBB leakage phenotypes present as hotspots of intense tracer accumulation rather than a homogenous distribution of tracer in the brain? Are specific vessel subtypes or brain regions more susceptible to leakage? If so, what is the biological basis for this susceptibility? Are these leakage hotspots static or is there dynamic loss and/or repair of BBB integrity?

HaloTrace’s capacity for multi-timepoint, longitudinal measurement could also enable researchers to relate transient leakage phenotypes to later outcomes. For example, one could test whether BBB leakage severity after a traumatic brain injury^28^ or during lesion formation in a multiple sclerosis model corresponds to worse cognitive outcomes in the days to weeks after the BBB leakage has resolved.^29,30^ HaloTrace could similarly be used to characterize the temporal dynamics of leakage onset in a mouse model or to describe the time course of successful BBB opening and closure during an intervention like focused ultrasound treatment.^31,32^

One of the most exciting potential applications of HaloTrace is *in vivo* fluorescence imaging. HaloTrace’s negligible signal in blood holds promise for visualizing BBB leakage dynamics in real time with *in vivo* imaging of mouse brain. During *in vivo* imaging, the intravascular signal from the high concentration of fluorescent tracers in the blood is often very high compared to all but the most severe leakage. It limits the ability of two-photon imaging to capture subtle leakage phenotypes. HaloTrace does not suffer from this issue so it could be applied to image more modest leakage phenotypes in real time. *In vivo* imaging with HaloTrace could also be used to answer long-standing questions about BBB permeability dynamics on a cellular level, such as comparing how the speed and extent of BBB leakage differs between a mouse model with a tight junction defect and a model with leakage through transcytosis. In addition to expanding our knowledge of fundamental BBB biology, understanding BBB permeability dynamics could inform dosage and administration paradigms for delivering CNS therapeutics across the BBB.

We look forward to the application of HaloTrace to the longstanding questions described above, given its unique spatiotemporal specificity and sequential measurements of BBB permeability. We envision that HaloTrace can be further customized and paired with emerging microscopy technologies to answer novel questions in BBB biology.

## METHODS

### Mouse models

Mice were housed in a standard vivarium with *ad libitum* access to food and water. All experiments were conducted in accordance with Harvard Medical School standards and local Institutional Animal Care and Use Committee (IACUC) protocol. C57BL/6J mice (JAX:000664) were acquired from Jackson Laboratory and maintained in-house. Cdh5:CreER^T2^ mice^33^ (MGI:3848982) and Ctnnb1^f/f^ mice^34^ (JAX:004152, MGI: J:67966) were maintained in-house on a C57BL/6 background and crossed to produce Cdh5:CreER^T2/+^;Ctnnb1^f/f^ and Ctnnb1^f/f^ littermate controls. The Cdh5:CreER^T2/+^ mouse is a PAC transgenic line generated with the CreER^T2^ recombinase gene under the control of a Cdh5 promoter sequence. Ctnnb1^f/f^ mice have loxP sites flanking exons 2-6 of the *Ctnnb1* gene.

### Tamoxifen induction of gene knockdown

Tamoxifen was administered to Cdh5:Cre^ERT2/+^;Ctnnb1^f/f^ and Ctnnb1^f/f^ littermate control mice to induce acute knockdown of the β-catenin gene in endothelial cells. Tamoxifen (Sigma Aldrich, T5648-1G) was dissolved in peanut oil (Fisher Scientific, #S25760) at a concentration of 20 mg/mL. Starting on day one, mice received five consecutive days of intraperitoneal injections at a dose of 0.1 mg tamoxifen/g bodyweight/day. Tamoxifen administration was timed so BBB leakage was measured on day eight, except for experiments where BBB leakage was measured consecutively on days eight and nine. This tamoxifen regimen and experimental timeline was designed to measure BBB integrity days prior to the seizure development and morbidity that has been observed at later timepoints in the Cdh5:Cre^ERT2/+^;Ctnnb1^f/f^ mouse.^9^

### Adeno-associated virus

Expression of HaloTag receptor in mice was achieved by systemic delivery of custom adeno-associated viruses (AAVs) produced by the Janelia Viral Tools Team or Boston Children’s Hospital Viral Core. All custom AAV expression cassettes were subcloned into a plasmid AAV backbone (a gift from Viviana Gradinaru, Addgene #104061)^35^ which included a Woodchuck Hepatitis Virus Posttranscriptional Regulatory Element (WPRE) and hGH poly-adenylation site. A plasmid map of the HaloTag receptor expression vector is included in **Supplementary Figure 1A**. The extracellular HaloTag receptor AAV includes a CAG promoter followed by a coding sequence containing the following elements: an Igĸ gene leader sequence,^21^ the HaloTag receptor,^17^ the transmembrane domain of PDGFR,^36^ mScarlet fluorescent protein,^23^ and the C-terminal endoplasmic reticulum (ER) export peptide from Kir2.1.^22^ For HaloTrace experiments in which two fluorescent ligands were injected, an equivalent HaloTag AAV with GFP replacing mScarlet was used to avoid fluorescent spectral overlap during imaging. All HaloTag AAVs were packaged in AAV2/PHP.eB^25^ to target diverse CNS cells including neurons, astrocytes, and other populations. HaloTag AAV was delivered at a titer of 2e11 viral genomes via tail vein injection three weeks prior to the BBB permeability assay measurement.

### Tracer injections

#### HaloTag ligand

To measure blood-brain barrier permeability, 100 nmol HaloTag ligand conjugated to the cell-impermeable fluorescent dye JF635i or JF549i (HHMI Janelia) was suspended in 100 µL 1X phosphate buffered saline solution (PBS, Sigma, #P4417) and injected into systemic circulation via the tail vein three weeks after AAV-HaloTag administration. HaloTag ligand was allowed to circulate for 0.75 or 24 hours prior to anesthesia induction for perfusion (noted in results section).

#### Sulfo-NHS-Biotin

To measure blood-brain barrier permeability, EZ Link Sulfo-*N*- Hydroxysulfosuccinimide-LC-Biotin (Thermo Scientific #21335, 556.59 g/mol) was freshly suspended in PBS at 0.2 mg/g bodyweight (along with 100 nmol JF549i HaloTag ligand) and injected into systemic circulation via the tail vein. The tracer cocktail was allowed to circulate for 0.75 h.

### Histology

#### Perfusion fixation and sample collection

For all mouse experiments, unless otherwise noted, a transcardial perfusion fixation protocol was performed to remove blood contents from the brain prior to histological analysis. Mice were deeply anesthetized with a solution of ketamine hydrochloride (Zoetis, #40027676, working concentration 10mg/mL) and xylazine (Akorn, #59399-110-20, working concentration 2 mg/mL) in PBS administered at 15 µL/g bodyweight. At 15 minutes, after confirming the absence of toe pinch response, the body cavity was exposed and the dorsal rib cage removed. After insertion of a needle into the left ventricle, the right atrium was snipped, and 25 mL of cold 4% paraformaldehyde in PBS was perfused at a rate of ∼8 mL/min using a peristaltic pump (Avantor, 70730-062). Immediately following perfusion, brains were carefully dissected and placed into 4% paraformaldehyde in PBS and postfixed at 4C overnight with gentle rocking.

#### Tissue sectioning

PFA-fixed brains were washed with PBS three times for 20 minutes each to remove residual fixative. Then, sagittal sections of 50 µm thickness were collected with a vibratome (Leica, VT1200S). Sections were directly mounted on slides (VWR, #48311-703) or collected in PBS for immunofluorescence staining on floating sections.

#### Immunofluorescence staining

Staining was performed on slide-mounted sections except for HaloTrace colocalization with CD31, which was performed with floating sections. Slide-mounted or floating sections were permeabilized in 0.5% PBST (PBS with 0.5% Triton-X100 (Sigma, T8787)) for 20 minutes, then incubated in blocking buffer (consisting of 0.1% PBST with 10% normal donkey serum (NDS, Jackson ImmunoResearch, 017-000-121)) for 1 hour at room temperature with gentle rocking. Sections were incubated with primary antibody in 2% NDS in PBST at 4°C overnight with gentle rocking. Sections were washed in 0.1% PBST three times for 15 minutes then incubated with secondary antibody in 2% NDS in 0.1% PBST for two hours at room temperature with gentle rocking. Sections were washed twice in 0.1% PBST and incubated with DAPI (Thermo Scientific, PI62247, working concentration 0.2 µg/mL) in 0.1% PBST to stain nuclei. Sections were washed a final time in PBS for 15 minutes. At this time, floating sections were mounted on coverslips. Finally, sections were sealed with coverslips (VWR, #48393-251) applied with Fluoromount-G (Electron Microscopy Sciences, #17984-25) for microscopy. The following primary antibody was used: Goat anti-CD31 (R&D Systems, #AF3628, 1:100). The following corresponding secondary antibody was used at a 1:500 dilution: Donkey anti-goat-AF488 (Jackson ImmunoResearch, #705-545-147). Sulfo-NHS-biotin was detected with AF647-conjugated streptavidin (Thermo Fisher, #S32357, 1:500) during the secondary antibody step.

### Human embryonic kidney (HEK) cell transfection and HaloTag ligand application

HEK293T cells (Clontech, #632273) were cultured in DMEM (Corning, #10-017-CM) supplemented with 10% fetal bovine serum (R&D Systems, #S11150) and 1% penicillin-streptomycin solution (Thermo Fisher, #15140122). To characterize extracellular HaloTag expression and ligand properties, cells were plated on poly-L-lysine (Sigma, # P4707)-coated coverslips (Electron Microscopy Sciences, #72230-01). Using the Lipofectamine 2000 reagent (Invitrogen, #11668019), cells were transfected with extracellular HaloTag plasmid (identical to the AAV vector plasmid) or a cytosolic HaloTag plasmid containing the coding sequence for HaloTag without membrane targeting motifs. Twenty-four hours after transfection, HaloTag ligand conjugated to either JF635i or JF549i (Promega, in alpha testing*) was introduced into the media at 150nM. After incubation with ligand for 0.5 hours (or 24 hours, when noted in the text), the cells were washed with media, washed with PBS, and fixed with 4% paraformaldehyde for 15 minutes. Cells were washed again with PBS and incubated with 0.2 µg/mL DAPI (Thermo Scientific, PI62247) in PBST 0.1% for 15 minutes before a final PBS wash. Coverslips were mounted on slides (VWR, #48311-703) with Fluoromount-G (Electron Microscopy Sciences, #17984-25) for confocal microscopy.

*Note: although the ligands used for cell culture were produced by Promega rather than Janelia, they are licensed by Janelia and have the same spectral properties as the Janelia-derived ligands.

### Microscopy

Confocal Microscopy was conducted with a Leica SP8 laser scanning confocal (10x, 0.4 N.A.; 40x, 1.10 N.A.; 63x, 1.4 N.A.). An Olympus VS200 Slide Scanner; (20x, 0.8 N.A.) was used to image a broad overview of AAV-HaloTag expression.

### Data analysis and statistics

#### Representative images, graphs and figures

Image processing was performed in FIJI (NIH).^37^ Graphs were created in Prism v10.0.3 (GraphPad Software). Figures were assembled in Adobe Illustrator.

#### Quantification of in vitro HaloTag data

Single plane confocal images at 63x magnification were analyzed in CellProfiler v.4.2.5^38^ using a custom pipeline. Whole cell ROIs were identified with automated threshold-based object identification, and mean intensity of HaloTag receptor (i.e., mScarlet fluorescence) and ligand fluorescence were calculated for each cell.

#### Quantification of HaloTrace BBB leakage index

Confocal image stacks acquired at 10x magnification were max intensity projected with FIJI. Multiple ROIs from each mouse were collected. Quantification was performed blinded to genotype in CellProfiler using a custom pipeline. Unadjusted max intensity projections were segmented with an object recognition algorithm to identify areas positive for HaloTag receptor and ligand, respectively. Total coverage area and mean intensity of the HaloTag receptor and ligand, as well as total cerebellum area, were calculated for each image.

#### Quantification of HaloTrace hotspot overlap and colocalization

Hotspot overlap analysis was performed blinded to genotype in CellProfiler. JF549i and JF635i-ligand-positive areas were identified as described above. The total area containing at least one ligand was calculated, as was the total “overlap” area in which both ligands were present. Hotspot colocalization analysis was performed in FIJI using the JACoP plugin pipeline to calculate the Mander’s correlation coefficients and Pearson’s correlation coefficients.^39^ For the Mander’s correlation coefficient, a standard threshold was applied to all images to segment ligand-containing areas from image background.

#### Statistics

Statistical analysis was performed in GraphPad Prism. Sample size was determined based on sample sizes in similar studies. Linear regressions describing the relationship between HaloTag receptor and ligand fluorescence intensity *in vitro* were calculated for each group. For two-way comparisons, nested t tests were performed, except for the analysis of *in vitro* ligand fluorescence intensity, where Welch’s t test was performed to account for different standard deviation between groups. For multiple comparisons, nested ANOVAs with Tukey’s correction were performed. For all analyses, significance was considered at p<0.05. Graphs were generated in Prism with standard error of the mean displayed as error bars for all data. Details of statistical tests for each experiment are included in figure legends.

## ACKNOWLEDGEMENTS

We thank our colleagues in the Gu lab for constructive and insightful comments on this manuscript; Gary Yellen and Jonathan Cohen for thoughtful feedback; the Lavis lab and the Open Chemistry team at Janelia for their generous gift of JF dyes; Ken Chan and Ben Deverman for consultation about AAV-PHP.eB; Juan Martinez-Francois and Gary Yellen for assisting with early pilot experiments; and Ralf Adams for the Cdh5:Cre^ERT2^ mouse line. Imaging, consultation and/or services were in part performed in the Neurobiology Imaging Facility at Harvard Medical School which is supported in part by the HMS/BCH Center for Neuroscience Research as part of an NINDS P30 Core Center grant (NS072030). This work was supported by the National Science Foundation Graduate Research Fellowship (H.Z.), the National Institute on Aging (T32AG000222) (H.Z.), and the National Institute of Neurological Disorders and Stroke (DP1-NS092473 and R35NS116820) (C.G.). C.G. is a Howard Hughes Medical Institute Investigator.

## AUTHOR CONTRIBUTIONS

H.Z. and C.G. conceived of the study, interpreted results, compiled figures and wrote the manuscript with input from all authors. H.Z., E.K. and P.M. performed the experiments and analyzed the data.

## LEAD CONTACT

Correspondence and material requests should be addressed to Chenghua Gu

---

**Supplementary Figure 1.**
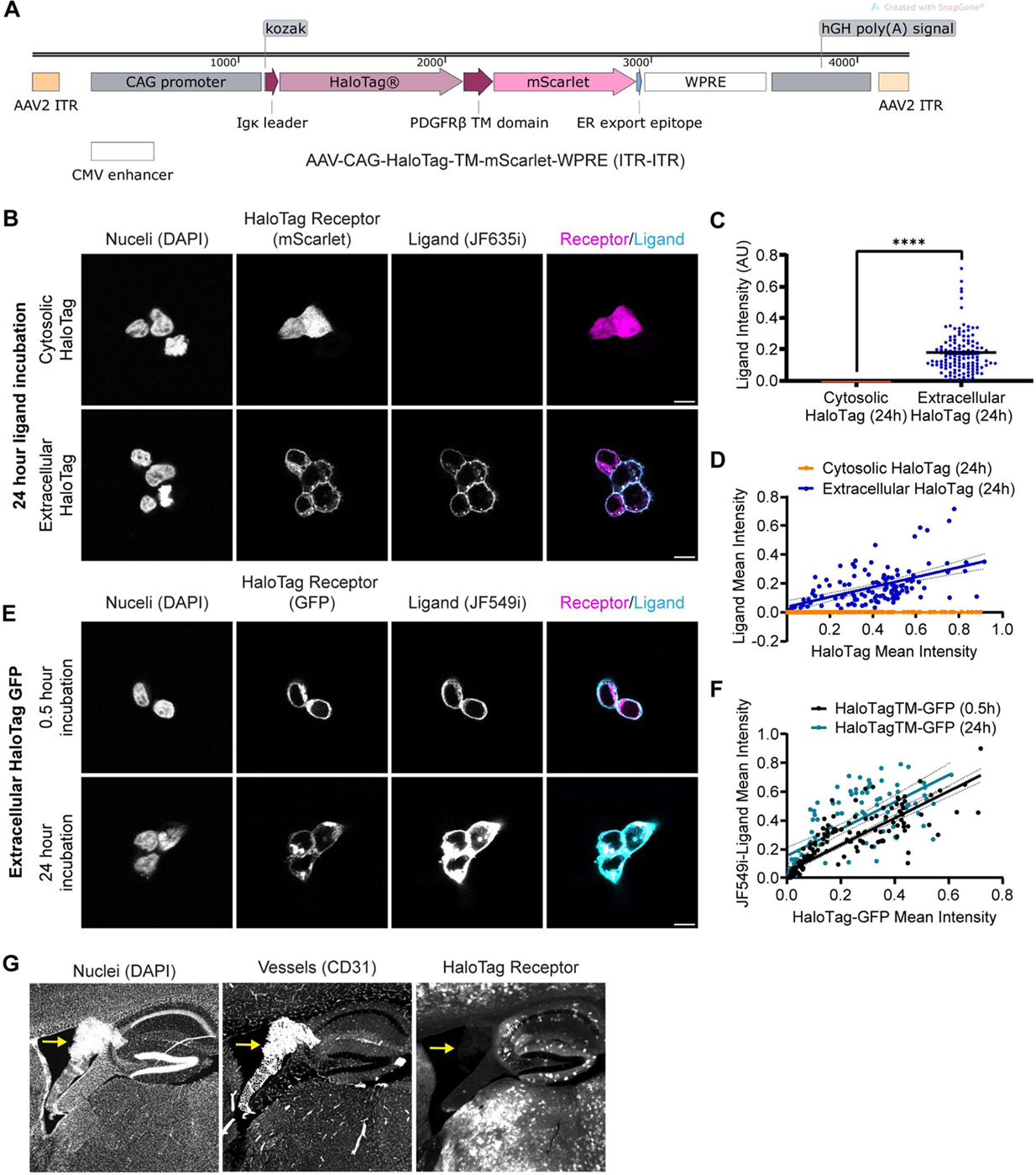
Further characterization of HaloTrace components. **(a)** Map of the AAV plasmid insert containing the CDS of extracellular membrane-targeted HaloTag fused to mScarlet. **(b)** 63x magnification confocal images of HEK cells transfected with cytosolic-targeted HaloTag-mScarlet or Extracellular HaloTagTM-mScarlet (magenta) incubated with JF635i-ligand (cyan) for 24 hours. Nuclei visualized with DAPI. Scale bar: 10 µm. **(c)** Quantification of JF635i ligand fluorescence intensity in extracellular HaloTag (blue) and cytosolic HaloTag (orange) cells. Data are mean ± SEM, extracellular n = 124, cytosolic n = 153 cells, ****p< 0.0001, Welch’s unpaired t test. **(d)** Relationship between HaloTag receptor expression (mScarlet fluorescence) and JF635i ligand fluorescence intensity in extracellular HaloTag and cytosolic HaloTag conditions. Data points represent single cells transfected with extracellular HaloTag (blue, n=124) or cytosolic HaloTag (orange, n=153). Linear regression for extracellular HaloTag: y=0.3449x+0.0367, r^2^=0.30. Linear regression for cytosolic HaloTag: y=0.0013x+ 0.0002, r^2^=0.66. **(e)** 63x magnification confocal images of HEK cells transfected with extracellular HaloTagTM-GFP (magenta) incubated with JF549i-ligand (cyan) for 0.5 or 24 hours. Nuclei visualized with DAPI. Scale bar: 10 µm. **(f)** Relationship between HaloTag receptor expression (mScarlet fluorescence) and JF635i ligand fluorescence intensity in extracellular HaloTag and cytosolic HaloTag conditions. Data points represent single cells incubated with ligand for 0.5h (black, n=186) or 24h (teal, n=91). Linear regression for 0.5h incubation condition: y=0.9323x+0.0431, r^2^=0.74. Linear regression for 24h incubation condition: y=0.9388x+ 0.1520, r^2^=0.50. **(g)** Slidescanner image of periventricular brain region of a C57Bl6 mouse injected with AAV-HaloTagTM-mScarlet. Nuclei, vessels, and HaloTagTM-mScarlet all shown in grayscale. Yellow arrow points to the lateral ventricle choroid plexus, which is not transduced.

**Supplementary Figure 2.**
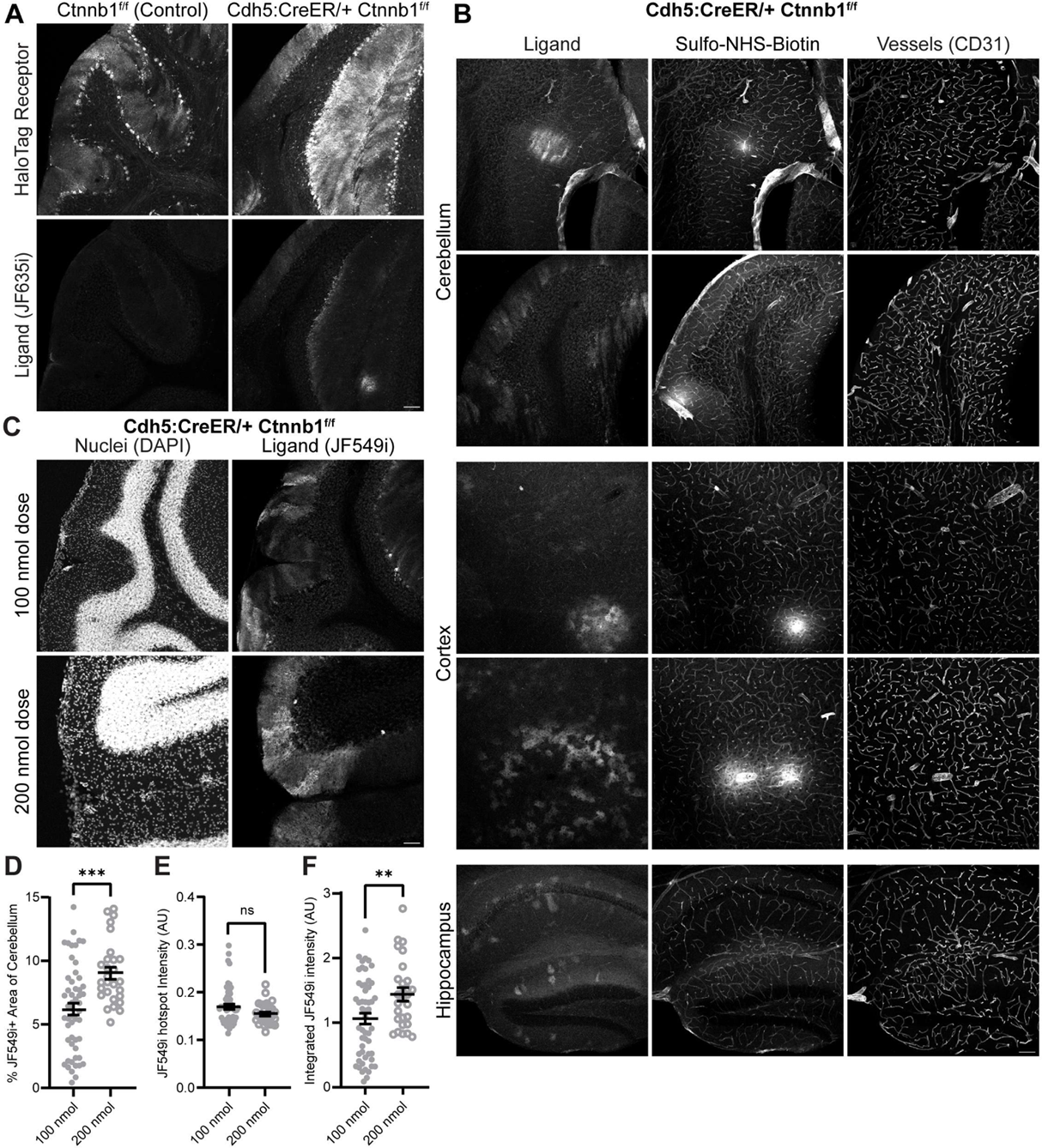
Further characterization of HaloTrace in Ctnnb1 knockdown. **(a)** Representative 10x magnification confocal images of HaloTrace permeability assay in Cdh5:CreER/+; Ctnnb1^f/f^ and control Ctnnb1^f/f^ cerebellums processed by dropfixation without transcardial perfusion. **(b)** Representative 10x magnification confocal images of Cdh5:CreER/+; Ctnnb1^f/f^ brains with BBB leakage assayed by HaloTrace and sulfo-NHS-biotin. Vessels visualized with anti-CD31 antibody. **(c)** Representative 10x magnification confocal images of ligand deposition in Cdh5:CreER/+; Ctnnb1^f/f^ cerebellum after injection of 100 nmol or 200 nmol JF549i ligand. Scale bar: 100µm. **(d)** Quantification of total JF549i-ligand containing area of cerebellum. *** indicates p=0.002, unpaired t test. **(e)** Quantification of JF549i+ area mean intensity. ns, nonsignificant, p=0.055, unpaired t test. **(f)** Quantification of integrated intensity of JF549+ positive area. ** indicates p=0.008, unpaired t test. For b-d, data points represent individual images from n=3 mice per dosage group. Filled gray circles: 100 nmol dose. Empty gray circles: 200 nmol dose. Data for the 100 nmol condition were also used in the analysis of Figure 4. For a,c,d, Scale bar: 100 µm.

**Supplementary Figure 3.**
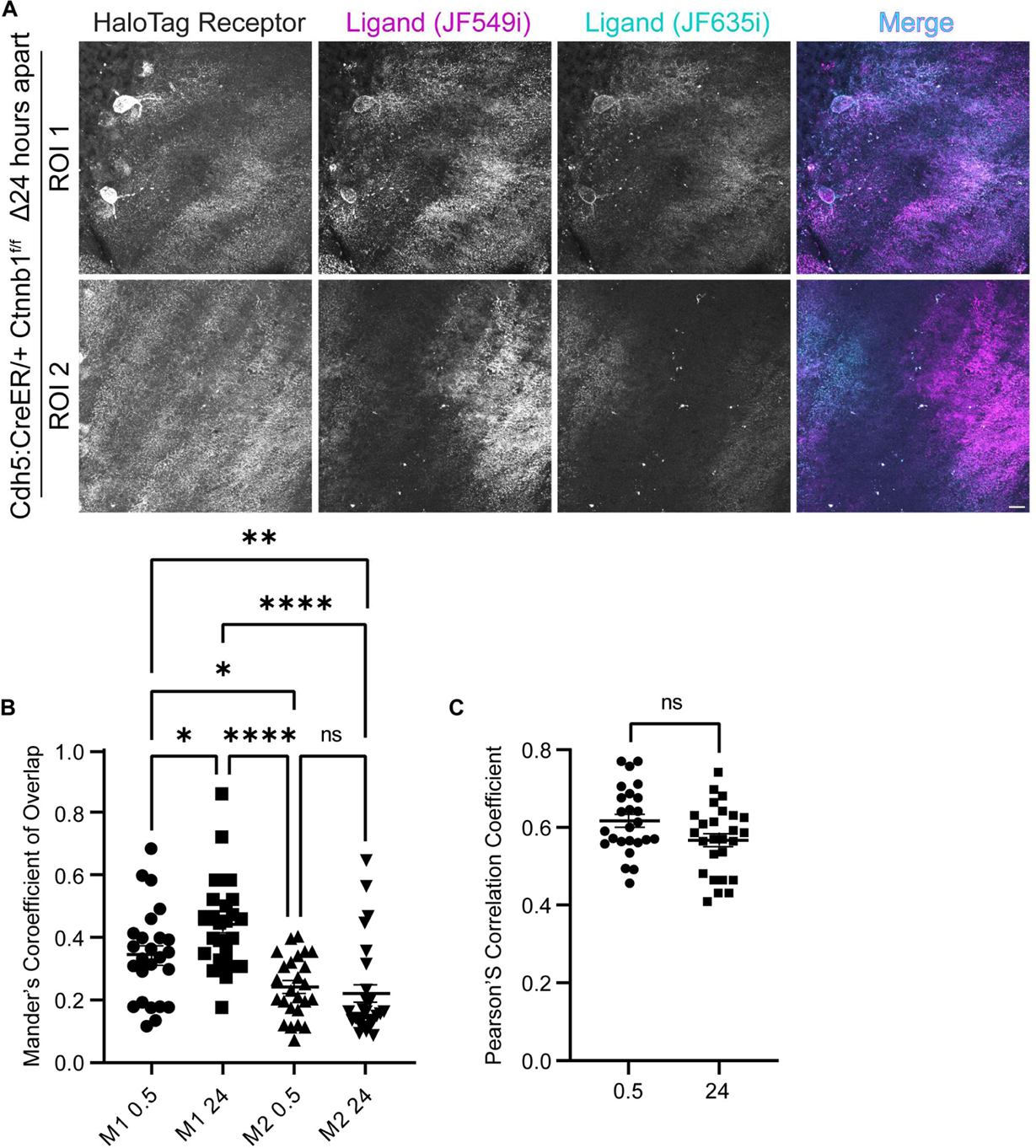
Intensity-based colocalization analyses of the pattern of JF635i- and JF549i-ligand. **(a)** 40x magnification confocal images of HaloTag receptor, JF549i-ligand (magenta in merge) and JF635i-ligand (cyan in merge) in Cdh5:CreER/+; Ctnnb1^f/f^ cerebellum. The top row shows an example of highly colocalized ligands whereas the bottom row shows an example of relatively distinct ligand hotspots. Scale bar: 20 µm. **(b)** Manders colocalization analysis. M1 corresponds to the fraction of JF549i overlapping JF635i in the Δ0.5h or Δ24h conditions; M2 corresponds to the fraction of JF635i overlapping JF549i. *p<0.05, ****p<0.0001, ns: nonsignificant, ANOVA with Tukey’s correction for multiple comparisons. **(c)** Pearson’s correlation coefficient as a measure of JF635i and JF549i-ligand overlap in the Δ0.5h or Δ24h conditions. ns: nonsignificant, student’s t test. p=0.055. For both analyses, n = 25 images in Δ0.5h group and n = 26 for the Δ24h group. Note that Pearson’s correlation coefficient can overestimate intensity correlation in samples with high background, like these, because it does not exclude background pixels from the analysis.

